# *In vivo* two-photon FLIM resolves photosynthetic properties of maize bundle sheath cells

**DOI:** 10.1101/2024.10.07.617075

**Authors:** Zhufeng Chen, Jing Li, Baichen Wang, Lijin Tian

**Author notes:** Lijin Tian **Email:**.

## Abstract

Maize (*Zea mays* L.) performs highly efficient C_4_ photosynthesis by dividing photosynthetic metabolism between mesophyll and bundle sheath cells. *In vivo* physiological measurements are indispensable for C_4_ photosynthesis research as any isolated cells or sectioned leaf often show interrupted and abnormal photosynthetic activities. Yet, direct *in vivo* observation regarding bundle sheath cells in the delicate anatomy of the C_4_ leaf is still challenging. In the current work, we used two-photon fluorescence-lifetime imaging microscopy (two-photon-FLIM) to access the photosynthetic properties of bundle sheath cells on intact maize leaves. The results provide spectroscopic evidence for the diminished total PSII activity in bundle sheath cells at its physiological level and show that the single PSIIs could undergo charge separation as causal. We also report an acetic acid-induced chlorophyll fluorescence quenching on intact maize leaves, which might be a physiological state related to the nonphotochemical quenching mechanism.

## Introduction

Photosynthesis is the primary energy input into the crop yield. Enhancing photosynthesis as an explicit breeding target is getting increasingly accepted, even though photosynthesis may have been subtly improved over the years during traditional breeding. Identifying the bottlenecks that limit photosynthetic efficiency is essential for achieving this breeding goal. Many strategies for improving photosynthesis have been explored, and they have been recently reviewed in ref. (Croce et al., 2024). However, as are often the cases, successful and unsuccessful examples have been reported when the same approach was applied to varied species, asking for costumed strategies for different crops. Clearly, species-specific phenotypes, e.g., within the leaf, should be considered for the next generation of breading. When spatial multi-omics technologies are available, photosynthetic traits on the cellular level should also be collected parallelly. Proper tools and methods that could probe the photosynthetic features on cell or sub-cellular levels *in vivo* are thus urgently demanded.

C_4_ plants are hallmarked by the capacity of photosynthetic CO_2_-concentrating mechanism, which relies on spatial differentiation of photosynthetic metabolism, usually between two types of leaf cells (Hatch, 1987). Of particular interest are the keystone model C_4_ plants like maize (*Zea mays* L.), with their Kranz anatomy composed of bundle sheath cells (BSCs) and mesophyll cells (MCs) considered to be a masterpiece of C_4_ metabolic differentiation. Various studies are dedicated to differentiating the two types of cells in terms of biochemical composition and physiological functions. Transcriptome, proteomic, and metabolome studies of purified mesophyll and bundle sheath cells provides an elaborative picture of cell metabolic division, suggesting reduced abundance of active photosystem II (PSII) and limited linear electron transport in BSC, while metabolic reactions consuming reduction equivalent are mainly located in MC (Majeran et al., 2008; Friso et al., 2010; Li et al., 2010; Majeran et al., 2010; Pick et al., 2011; Ponnala et al., 2014; Tausta et al., 2014). Biochemical as well as physiological analyses on sliced leaves or isolated leaf cells roughly show a lower PSII activity in bundle sheath cells than in mesophyll cells (Shibata et al., 2013; Chiba and Shibata, 2019; Hernandez-Prieto et al., 2019; Jackowski et al., 2022), which is further supported by results base on simulated metabolic studies (Bellasio and Griffiths, 2013; Wang et al., 2014; Ludwig, 2016; Yin and Struik, 2018). Even so, physiological-level evidence about the actual PSII activity in BSC remains lacking. Restricted by the complex structure of C_4_ leaf anatomy, an *in vivo* chlorophyll fluorescence measurement of bundle sheath cells is challenging.

Two-photon fluorescence-lifetime imaging microscopy (two-photon-FLIM), as a non-invasive technique with deep tissue penetration, provides the possibility to probe BSC *in vivo*, and the concept of using two-photon FLIM to study C_4_ BSC was first validated on C_4_ plant *Miscanthus × giganteus* (Iermak et al., 2016). Also, FLIM, as an extension of conventional time-resolved fluorescence techniques, is increasingly used for *in vivo* research on photosynthetic light-harvesting regulatory mechanisms such as state transitions (Wientjes et al., 2017; Bhatti et al., 2021; Verhoeven et al., 2023). This study uses two-photon-FLIM to obtain deep-tissue time-resolved fluorescence images of intact maize leaves with fluorescence decay profiles of both BSC and MC. The differences in photosynthetic light-harvesting properties between MC and BSC are explored under a physiological and artificially induced quenching state.

## Results

### Visualize and distinguish the fluorescence decay kinetics of bundle sheath cells in the maize leaf

The bundle sheath cells (BSCs) are located inside the leaf about 50 μm from the leaf surface (starting with the stomatal guard cells; see Figure 1a). At such a depth, the spatial resolution of imaging decreases sharply due to intense light scattering from the upper mesophyll cells (MCs, Figure 1b). To confirm the location of the BSC, the image of the cell profile in the same field of view was additionally obtained (Figure 1c, see method) and merged with the chlorophyll fluorescence image (Figure 1d). With the location and shape of BSC confirmed, only chlorophyll fluorescence images were considered in subsequent experiments. While the individual chloroplasts inside the BSC are hard to distinguish, each elongated BSC can be identified. Due to the limited axial resolution of the two-photon microscope, the out-of-focus fluorescence of the upper MC chloroplast might superimpose on the region corresponding to BSC, resulting in distortion in the kinetics. Therefore, the depth of the imaging field was carefully controlled to ensure that the fluorescence signals from BSC were not contaminated by MC fluorescence.

**Figure 1.**
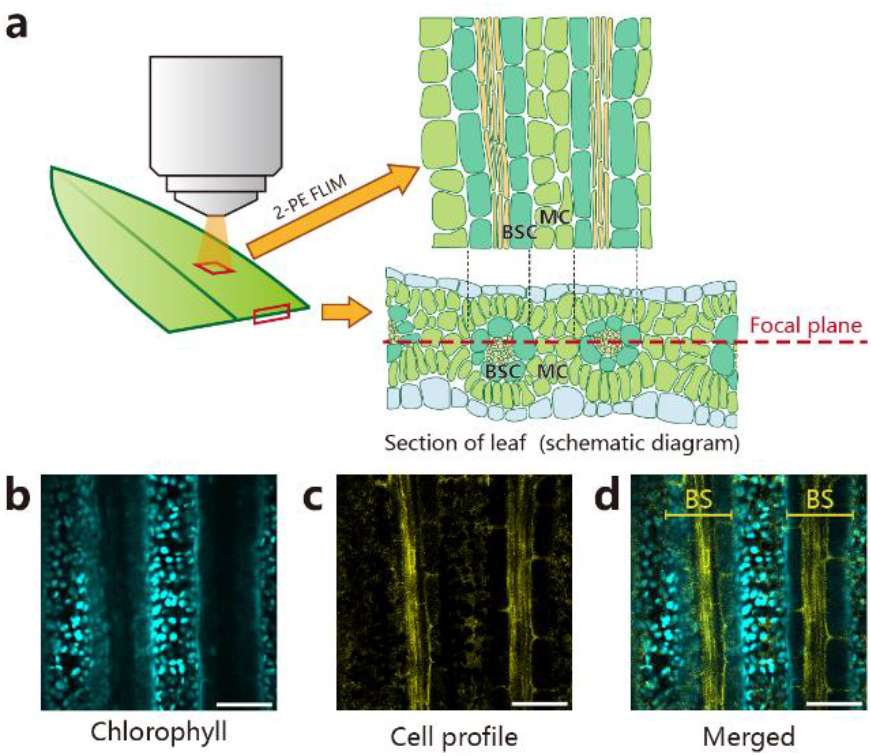
(a) Schematic diagrams of the leaf section parallel (top right) and vertical (bottom right) to the vein demonstrate the distribution of mesophyll cell (MC) and bundle sheath cell (BSC). The focal plane of the microscope is parallel to the leaf surface (red dotted line). (b) Steady-state chlorophyll fluorescence image of maize leaf detected at 655-725 nm. (c) Cell profile image of maize leaf bundle sheath detected at 500-550 nm, mainly derived from the second harmonic signal of the cell wall. For clear identification of the cell profile, the chlorophyll fluorescence signal is additionally acquired at 570-620 nm and then subtracted from the image. (d) A merged image of the cell profile and chlorophyll fluorescence with the bundle sheath (BS) position is highlighted. The scale bar is 50 μm.

### Assess FLIM data quality using the phasor approach

One of the major challenges in time-resolved fluorescence measurement is nonlinear effects (e.g., singlet-singlet (S-S) or singlet-triplet (S-T) annihilation processes between excited Chls) arising from high excitation density, especially for large pigments-binding complex of PSII in leaves. When the plane of focus goes deep into the leaf, the uneven distribution of upper-layer chloroplasts generates a dramatic horizontal variation of excitation density, leading to heterogeneous annihilation kinetics. The spatial heterogeneity of fluorescence kinetic originating from cytological diversity is therefore obfuscated by the heterogeneity of annihilation kinetics.

Instead of examining the fluorescence decay profiles by each pixel, we took advantage of the phasor approach (Digman et al., 2008) to quickly recognize the fluorescence kinetics pattern of the whole image and screen the suitable (annihilation-free) measuring conditions. The principle of Phasor-plot based segmentation is described in ref. (Ranjit et al., 2018) Briefly, the normalized Fourier sine and cosine transforms at a particular frequency of the photon histogram curve, namely S and G, are mapped to a two-dimensional space, the phasor space. Each data point in the phasor plot corresponds to a pixel of the FLIM image. Thus, the distribution of points in the phasor plot visually shows the lifetime characteristics of the sample. Figure S1 gives an example of how the phasor approach helps to assess the effect of annihilation on FLIM data quickly. With the increase of excitation power, the cluster of points in the phasor plot gradually shows a tail. The tail disappears when excitation power is turned low again, implying that it does not result from sample damage. This tail is an evident characteristic of annihilation kinetics distorting fluorescence lifetime measurement, which we tried to avoid during the measurements.

For the regions corresponding to MCs, heterogeneity occurs even at very low excitation power (Figure 2; see Figure S1). Potentially because the MCs between the two bundles are less shaded by the upper layer chloroplasts, subject to higher excitation density. In contrast, the BSCs in the same depth are more shielded and receive lower excitation density, less affected by annihilation kinetics (see Figure 1a for a schematic). For comparison, fluorescence decay profiles were additionally obtained from images focused on the upper layer of the leaf, targeting MCs only, using much lower excitation power (Figure 3). It can be seen in the phasor plot that the points of mesophyll cells on the leaf surface are less affected by annihilation (Figure S2).

**Figure 2.**
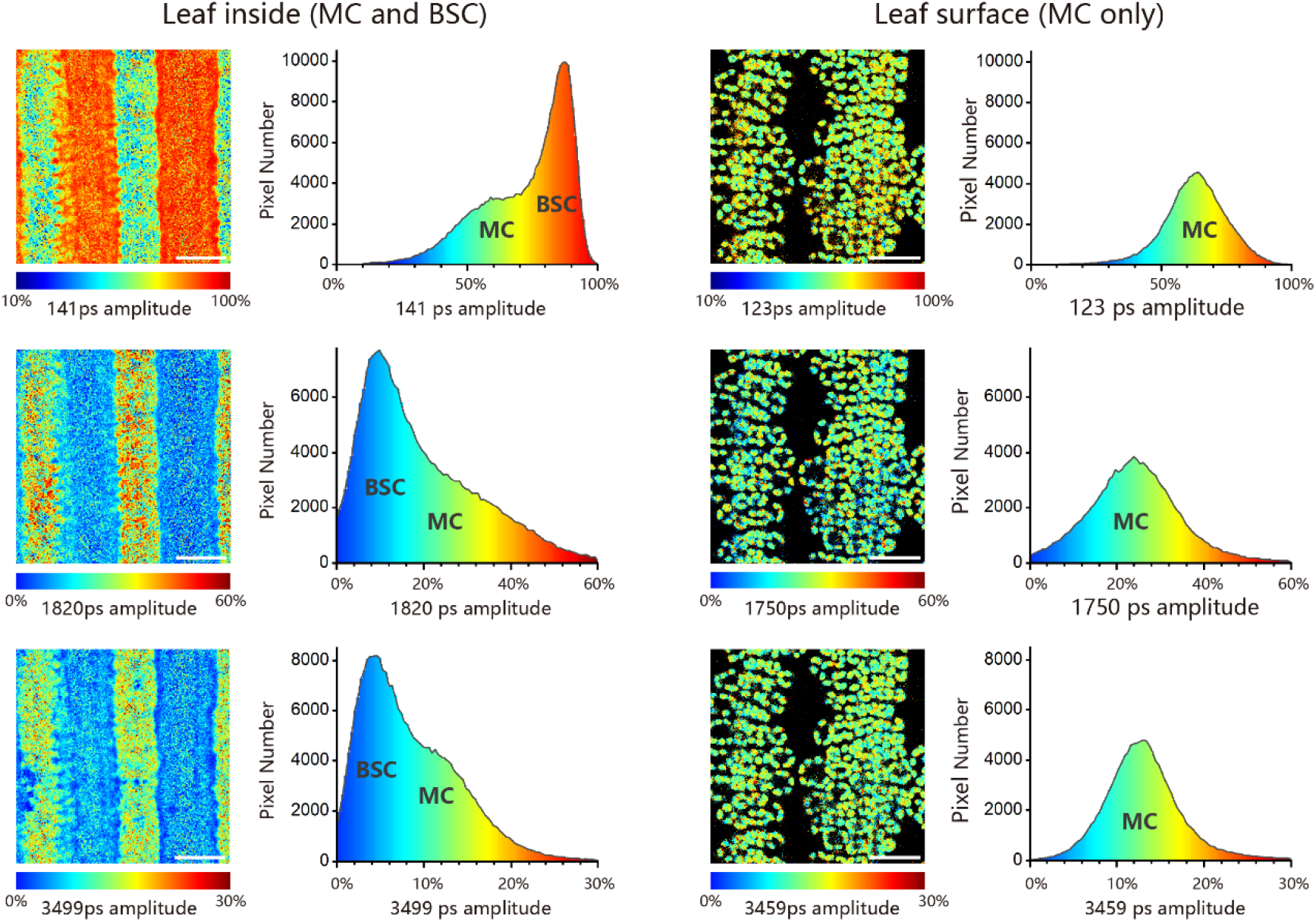
Fitting result to maize leaf FLIM data (view of leaf inside and leaf surface) with a three-exponential decay model. The pseudo-color images (left) show the amplitudes of lifetime components in each pixel, while the histograms (right) summarize the number of pixels with the same amplitude value. In all samples, PSII RCs are chemically closed by DCMU and hydroxylamine treatment. The scale bar is 50 μm.

**Figure 3.**
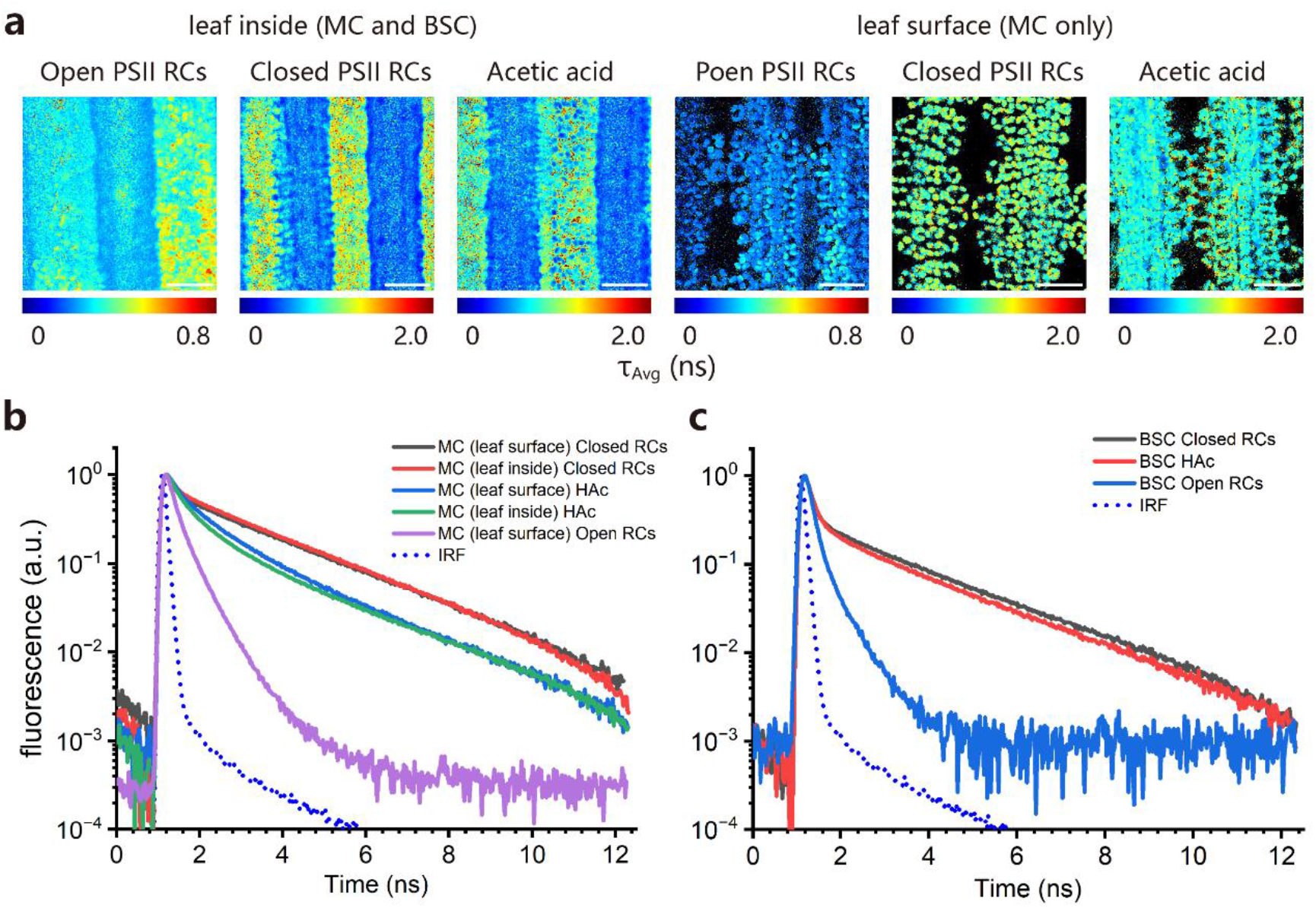
(a) Average lifetime images of maize leaf (focus inside the leaf and on the leaf surface) under open PSII RCs (leaves infiltrated with water), closed PSII RCs (leaves treated with DCMU and hydroxylamine before infiltrated with water) and acetic acid (HAc) treatment (leaves treated with DCMU and hydroxylamine before infiltrated with acetic acid) conditions. The scale bar is 50 μm. (b) Fluorescence decay profiles of mesophyll cells (MC, on the leaf surface and inside the leaf) under open PSII RCs, closed PSII RCs, and acetic acid treatment conditions. (c) Fluorescence decay profiles of bundle sheath cells (BSC) under open PSII RCs, closed PSII RCs, and acetic acid treatment conditions.

### Photosynthetic characteristics of mesophyll and bundle sheath cells

One of the major characteristics of maize leaf BSCs is their reduced PSII/PSI ratio, which provides an intuitive way to discriminate them from MCs since the fluorescence kinetics of PSI and PSII can be separated *in vivo* by using time-resolved fluorescence techniques (van Oort et al., 2010; Chukhutsina et al., 2015; Tian et al., 2017). With reduced PSII activity (Shibata et al., 2013; Chiba and Shibata, 2019; Hernandez-Prieto et al., 2019; Jackowski et al., 2022), BSC show a faster fluorescence lifetime than MC, particularly when PSII reaction centers (RCs) are photochemically closed (Q_A_ reduced). The MC and BSC signals can thus be segmented in the phasor plot based on their diverse fluorescence kinetics (Figure S2).

With PSII RCs chemically closed (see methods), the fluorescence decay profiles of MC and BSC can be fitted with a three-exponential decay model, giving three lifetime components similar in both types of cells, including a fast lifetime component of 120-140 ps and two slow lifetime components of ∼1800 ps and ∼3500 ps, respectively (Table 1). The ∼1800 ps and ∼3500 ps components are assigned to PSII with closed RCs. The 120-140 ps component is mainly contributed by PSI, also containing a small fraction of PSII faster component that cannot be distinguished due to the temporal resolution of the experimental setup (IRF ∼160 ps FWHM), as evidenced by the slight increase of its amplitude in the case of open PSII RCs (Q_A_ oxidized). With this small fraction of PSII fast kinetics ignored, the contribution of PSII can be estimated through the slow lifetime component amplitudes, giving the PSII/(PSII+ PSI) activity ratio of 0.45 and 0.17 in MC and BSC, respectively.

**Table 1.** Fitting result to fluorescence decay kinetics of maize leaf mesophyll cell (MC) and bundle sheath cell (BSC) under open PSII RCs (leaves infiltrated with water), closed PSII RCs (leaves treated with DCMU and hydroxylamine before infiltrated with water) and acetic acid (HAc) treatment (leaves treated with DCMU and hydroxylamine before infiltrated with acetic acid) conditions.

| Open PSII RCs |  |  |  | Closed PSII RCs |  |  |  | HAc treatment with closed PSII RCs |  |  |  |
| --- | --- | --- | --- | --- | --- | --- | --- | --- | --- | --- | --- |
| BSC |  | MC |  | BSC |  | MC |  | BSC |  | MC |  |
| $\tau_i$, ps | $a_i$, % | $\tau_i$, ps | $a_i$, % | $\tau_i$, ps | $a_i$, % | $\tau_i$, ps | $a_i$, % | $\tau_i$, ps | $a_i$, % | $\tau_i$, ps | $a_i$, % |
| 141 | 90.4 | 147 | 71.9 | 140 | 83.4 | 140 | 54.7 | 141 | 83.4 | 150 | 48.2 |
| 468 | 9.5 | 470 | 28.0 | 1817 | 11.7 | 1769 | 31.6 | 1371 | 10.9 | 573 | 32.8 |
| 18779 | 0.1 | 12212 | 0.1 | 3529 | 4.8 | 3428 | 13.7 | 3401 | 5.7 | 1727 | 16.6 |
|  |  |  |  |  |  |  |  |  |  | 5655 | 2.4 |
| $\tau_{Avg}$, ps | 196 | $\tau_{Avg}$, ps | 244 | $\tau_{Avg}$, ps | 501 | $\tau_{Avg}$, ps | 1104 | $\tau_{Avg}$, ps | 462 | $\tau_{Avg}$, ps | 681 |

### Chlorophyll fluorescence quenching induced by acetic acid on maize leaves

It was previously revealed in *Chlamydomonas* that photoprotective energy-dependent quenching of chlorophyll fluorescence (also known as qE) could be induced by artificially reducing lumen pH with weak acids like acetic acid (Pang et al., 2023). Here, we test the effectiveness of this method on maize leaves. As shown in Figure 3, with PSII RCs chemically closed, acetic acid treatment induces fluorescence quenching in both MC and BSC, which is more pronounced in MCs. It is also noted that MCs inside the leaf are slightly more affected by acetic acid than the leaf surface, presumably because the acetic acid enters the leaf sample along the vessel and preferentially affects MCs adjacent to the vascular bundle. The retention of slow lifetime components and the differences between MC and BSC suggest that acetic acid-treated samples may represent a physiological state, probably related to photoprotective quenching of PSII fluorescence.

The quenching capacity, calculated as (τ_Avg,uq_ − τ_Avg,q_)/τAvg,q according to the average lifetimes under unquench (τAvg,uq) and quench (τAvg,q) state induced by acetic acid, is much higher in MC than in BSC of 0.62 and 0.08, respectively. In addition, assuming that the difference of slow lifetime components (τ_slow_ = ∑ α_i_ · τ_i_ / ∑ α_i_, i for >150 ps components) before and after acetic acid treatment (in the case of PSII RCs chemically closed) represents the quenching of physiologically active PSII, the PSII quenching capacity is calculated as (τ_slow,uq_ − τ_slow,q_)/τ_slow,q_, giving the value of 0.93 and 0.12 for MC and BSC, respectively. The two calculations above provide an estimated range for the quenching capacity of both types of cells, which is ∼7 times higher in MC than BSC under current experimental conditions.

## Discussion

### Noninvasive time-resolved fluorescence measurements shed more light on bundle sheath cells

One well-known difficulty in leaf chlorophyll fluorescence measuring is its extreme susceptibility to mechanical injury, which complicates almost all results involving direct sample dissecting. In practice, cutting the leaves leads to an immediate quenching of chlorophyll fluorescence adjacent to the incision, manifested by the decreased intensity and shortened fluorescence lifetime (see Figure S3 for an example of an incision changing the chlorophyll fluorescence). This might cause the low Fv/Fm, ϕPSII, and NPQ values obtained from sliced leaves compared to intact leaves (Liu et al., 2022). Although the mechanical injury-induced quenching of chlorophyll fluorescence seems commonplace, how it happens remains uncharacterized. Given that various biotic or abiotic stresses tend to decrease chlorophyll fluorescence, its underlying mechanism could be an interesting scientific question for the future. In the current work, we bypassed this difficulty by utilizing two-photon microscopy with appropriate sample penetration. Combining intensity-independent fluorescence lifetime measurement, we calculated the energy distribution between the two photosystems, confirming the light-harvesting functional differentiation of maize MC and BSC in their functional states *in vivo*.

In addition to FLIM, there are mature protocols for time-resolved fluorescence measurements on intact leaves, including the so-called fluorescence lifetime snapshots (Sylak-Glassman et al., 2014; Sylak-Glassman et al., 2016) and Lissajous scanner method (Farooq et al., 2018; Chukhutsina et al., 2019; Chukhutsina et al., 2020). The former focuses on the dynamic change of fluorescence lifetime over long time scales (e.g., tens of minutes for induction and relaxation of non-photochemical quenching), while the latter focuses on multispectral fluorescence data acquisition under specific physiological states. The ∼1 ns average lifetime derived from the current work agrees with the above research, regardless of the different experimental setups and diverse ways to maintain closed PSII RCs. On the other hand, the temporal resolution of the experimental setup in the current work is inadequate for distinguishing the PSII fast-lifetime component from PSI kinetics (see above), which restricts the potential for the fine distinction of PSII/PSI fluorescence. Instead, the spatial variation of the PSII/PSI ratio, especially the difference between BSC and MC, is the primary concern of the current work.

### PSII activity in bundle sheath cells

The enhancement of cyclic electron flow (CEF) around PSI in BSC is considered to be critical for NADP-ME type C4 plants, including maize, as it provides the additional ATP required for C_4_ metabolism (Ishikawa et al., 2016). CEF mediated by the chloroplast NADH dehydrogenase-like complex (NDH) is the prime candidate for the presumptive C_4_-supporting CEF pathway, as evidenced by the substantial decrease in CO_2_ assimilation capacity in mutants deficient in NDH function among NADP-ME type C4 plants (Peterson et al., 2016; Ogawa et al., 2023; Ermakova et al., 2024; Zhang et al., 2024). This accounts for the high PSI activity observed in maize BSC here. On the other hand, our understanding of PSII activity in BSC remained fragmentary. Proteomic data suggests the deficiency of PSII in maize BSC, as evidenced by the down-regulation of several subunits of PSII core and oxygen-evolving complex (OEC) (Majeran et al., 2008). Thylakoid complexes analysis by Blue Native PAGE also reveals distinct PSII composition between MC and BSC despite the diversity across different studies (Hernandez-Prieto et al., 2019; Liu et al., 2022; Sárvári et al., 2022). The current results indicate that at least part of the PSII in BSC is fully physiologically active, as the slow lifetime components representing PSII are similar in BSC and MC with either open or closed RCs (Table 1).

So, is there a “perfect” PSII/PSI ratio for C_4_ BSC? To the best of our knowledge, very little is known. Modeling studies suggest that the flexibility of metabolite flux between MC and BSC can compensate for the imbalance of PSII/PSI excitation in two types of cells (Bellasio and Griffiths, 2013). Suppose the flexibility of C4 metabolite flux provides scope for variation for PSII/PSI activity. In that case, it is possible that the actual PSII/PSI activity in BSC is dynamically modified by factors like growth stage (Romanowska et al., 2006) and light conditions (Zienkiewicz et al., 2015; Rogowski et al., 2019). Correlational studies could be facilitated by the FLIM method described here.

### The acetic acid-induced chlorophyll fluorescence quenching may represent a physiological state

The acidification-induced chlorophyll fluorescence quenching in light-harvesting complexes under *in vitro* conditions has been reported and rationally considered as a simulation of the widespread photoprotective energy dissipation in higher plants, which is triggered by thylakoid proton motive force and lumen acidity (Petrou et al., 2014; Schlau-Cohen et al., 2015). With pH decreased, the major light-harvesting complex of PSII (LHCII) could allosterically transform to a photoprotective energy-quenching state, with the overall process fine-tuned by factors like PsbS protein and zeaxanthin (Johnson and Ruban, 2009; Nicol and Croce, 2021; Ruan et al., 2023). Starting from this model, the much smaller quenching capacity in BSC than in MC demonstrated here could be regarded as reduced sensitivity to overexcitation of PSII in BSC. Indeed, BSC surrounded by MC might experience less extreme light intensity, thus reducing the need for photoprotection. Moreover, PSII in BSC may play a relatively minor role compared with PSI and, therefore, be less regulated. Nevertheless, the physiological properties of the acetic acid-induced fluorescence quenching in maize leaves revealed here remain to be determined. Quantitative studies based on mutants related to light-harvesting regulation are needed, where the FLIM approach presented here will be advantageous for *in vivo* research under their physiological states.

## Materials and Methods

### Plant growth conditions and sample preparation

Maize plants, *Zea mays* L. line B73, were cultivated indoors in commercial potting soil at temperatures around 25 °C at a light intensity of 200 μE m^−2^ s^−1^ under plant culture LED light with a 14-h light period. The fourth or fifth leaves on 4- to 6-week-old seedlings were used. Leaves were taken during the light period to ensure enough starch accumulated, which provides resistance against infiltration and vacuum treatments. Before the measurement, leaves were infiltrated with 200 μM DCMU (3-(3’,4’-dichlorophenyl)-1,1-dimethylurea) and 20 mM hydroxylamine at room temperature in darkness for 2 hours (without detaching them from the plant). With this treatment, the PSII RCs were chemically closed while minimal impact was exerted on the leaves. For microscopic slides, leaves were cut into 5×10 mm pieces and then vacuum infiltrated with water inside a syringe. The water fills the gaps inside the leaf, reducing scattering and improving the imaging. For acid treatment, 50 mM acetic acid instead of water was used in vacuum infiltration.

### Two-photon fluorescence lifetime imaging and steady-state fluorescence imaging

Two-photon fluorescence lifetime imaging measurements were performed on inverted confocal microscopy (A1, Nikon). For the excitation of chlorophyll, 800 nm femtosecond laser (Chameleon Vision, Coherent, 80 MHz, 700 -1060 nm, 140 fs) was used. After passing through an attenuator, the fs laser beam was coupled into a laser confocal scanning system, deflected the beam while preserving the laser focus position in the focal plane of the objective (60×/0.95 NA, Nikon). The emitted fluorescence light was collected using the same objective as the one used for the excitation and sample scan. The short-pass filter (BrightLine multiphoton filter, 680 nm/SP, Semrock) was used to block the excitation fs laser light in the emission path. A band-pass filter (BrightLine 690/70 nm, Semrock) rejected the scattered excitation light. Finally, the emission light was sent into a single photon sensitive detector (PMA Hybrid 40, PicoQuant) based on a fast hybrid photomultiplier tube (R10467 from Hamamatsu) with a Peltier cooler to reduce the dark count rate. The output signal of the photon detector was recorded using a TCSPC system (HydraHarp 260, PicoQuant), which was synchronized with the triggering signal from the excitation laser. Data was acquired using commercial software from Picoquant (SymPhoTime 64, PicoQuant), which controlled both the TCSPC and the scanner systems. Instrument response function (IRF) was recorded with pinacyanol iodide in methanol, giving an IRF width of ∼160 ps FWHM. FLIM image was recorded within 15 min at a count rate of 10,000 counts/s (focus on the leaf surface) and 15,000 counts/s (focus deep into the leaf). The image size was 212 µm × 212 µm with 512 × 512 pixels. A steady-state fluorescence image was acquired with the same confocal microscopy. The image of the cell profile was detected at 500-550 nm, mainly derived from the second harmonic signal of the cell wall. The chlorophyll fluorescence signal was acquired at 570-620 nm for precise identification of the cell profile and then subtracted from the image.

### FLIM data processing

FLIM data were analyzed using the FLIMfit software tool developed at Imperial College London (Warren et al., 2013). A binning of 3×3 (leaf inside data) and 5×5 (leaf surface data) was used to improve the signal-to-noise ratio. Fluorescence lifetime was fitted following the ‘global binning’ approach. The decay profile data were integrated across the image to give a single decay profile and fitted to a sum of 3 exponentials 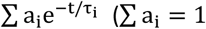, i runs from 1 to 3), convoluted with the instrument response function (IRF). The produced lifetime components were then fixed across the image, and the contributions of each component were determined by fitting them pixel-wise. Average lifetimes were calculated as τ_Avg_ = ∑ a_i_τ_i_. Phasor-plot based segmentation was carried out with FLIMfit software tool. The principle of Phasor-plot based segmentation is described in ref. (Ranjit et al., 2018)

## Acknowledgments and Funding

The authors thank Qingtao Lu, Dongyang Liu, and Peng Tian for their valuable comments on the manuscript, and Chunyan Zhang and Yan Yin for their excellent technical support. This research was supported by the Strategic Priority Research Program of the Chinese Academy of Sciences, Grant NO.XDA 26030201; and by the National Natural Science Foundation of China (No.31970381).

## Author Contributions

Z.C., B.W., and L.T. designed research; Z.C. and J.L. performed research; Z.C. analyzed data; and Z.C., J.L., B.W., and L.T. wrote the paper.

